# Unraveling three-dimensional chromatin structural dynamics during spermatogonial differentiation

**DOI:** 10.1101/2021.07.26.453830

**Authors:** Yi Zheng, Lingkai Zhang, Long Jin, Pengfei Zhang, Fuyuan Li, Ming Guo, Qiang Gao, Yao Zeng, Mingzhou Li, Wenxian Zeng

**Affiliations:** Key Laboratory for Animal Genetics, Breeding and Reproduction of Shaanxi Province, College of Animal Science and Technology, Northwest A&F University, Yangling, Shaanxi 712100, China; Institute of Animal Genetics and Breeding, College of Animal Science and Technology, Sichuan Agricultural University, Chengdu, Sichuan 611130, China

**Keywords:** Spermatogonia, 3D chromatin, A/B compartment, topologically associating domain, promoter-enhancer interaction

## Abstract

Spermatogonial stem cells (SSCs) are able to undergo self-renewal and differentiation. Unlike the self-renewal that replenishes the SSC and progenitor pool, the differentiation is an irreversible process committed to meiosis. While the preparations for meiotic events in differentiating spermatogonia (Di-SG) are likely to be accompanied by alterations in chromatin structure, the three-dimensional (3D) chromatin architectural difference between SSCs and Di-SG, and the higher-order chromatin dynamics during spermatogonial differentiation, have not been systematically investigated. Here, we performed *in situ* high-throughput chromosome conformation capture (Hi-C), RNA-sequencing (RNA-seq) and chromatin immunoprecipitation-sequencing (ChIP-seq) analyses on porcine undifferentiated spermatogonia (Un-SG, which consist of SSCs and progenitors) and Di-SG. By integrating and analyzing these data, we identified that Di-SG exhibited increased disorder but weakened compartmentalization and topologically associating domains (TADs) in comparison with Un-SG, suggesting that diminished higher-order chromatin architecture in meiotic cells, as shown by recent reports, is preprogramed in Di-SG. Our data also revealed that A/B compartments and TADs were related to dynamic gene expression during spermatogonial differentiation. We further unraveled the contribution of promoter-enhancer interactions (PEIs) to pre-meiotic transcriptional regulation, which has not been accomplished in previous studies due to limited cell input and resolution. Together, our study uncovered the 3D chromatin structure of SSCs/progenitors and Di-SG, as well as the interplay between higher-order chromatin architecture and dynamic gene expression during spermatogonial differentiation, providing novel insights into the mechanisms for SSC self-renewal and differentiation and having implications for diagnosis and treatment of male sub-/infertility.

## Introduction

With every heartbeat a man produces over 1000 spermatozoa, each in theory capable of generating a new-born child (Johnson, Petty, & Neaves, 1980). The highly efficient production of spermatozoa is reliant on spermatogenesis, an intricate process occurring in the testis during which the primitive spermatogenic cells, i.e., spermatogonial stem cells (SSCs), develop into mature spermatozoa (Jan et al., 2012). SSCs are the only adult stem cells in males with the ability to transmit genetic information to the next generation, and thus they have a series of desirable attributes, with some shared by other stem cell categories. First, they strike a balance between self-renewal and differentiation to preclude exhaustion while simultaneously safeguard the ongoing production of gametes. Second, being located at a specific place in the mammalian testis called “niche”, SSCs are orchestrated by a host of intrinsic and extrinsic factors with well-defined roles in SSC fate determination and behaviors (de Rooij, 2017; Makela & Hobbs, 2019). Third, SSCs are capable of relocating to the basement membrane and reestablishing donor-derived spermatogenesis after transplantation into the allogenic recipient testis, being an appealing target for treatment of male infertility (Kubota & Brinster, 2018; Mulder et al., 2016).

Despite the crucial roles of SSCs in maintenance of male fertility, distinct models regarding the SSC property and cellular hierarchy have been proposed. Specifically, while traditional models, which are principally based on histological observations, propose that only the most primitive undifferentiated spermatogonia (Un-SG), i.e., single spermatogonia (A_s_), have stem cell characteristics (de Rooij, 2017), most data from later studies are more in favor of a dynamic stem cell model illustrating context-dependent and plastic stemness (Lord & Oatley, 2017; Makela & Hobbs, 2019). Nonetheless, it has generally been accepted that stem cell potential is typically limited to a rare subpopulation of Un-SG. Intriguingly, the recent boom of studies employing single-cell RNA-sequencing (RNA-seq) methodology have uncovered the remarkable heterogeneity of SSCs (Suzuki, Diaz, & Hermann, 2019; Tan & Wilkinson, 2019, 2020), and with other omics approaches, the transcriptome, metabolome, DNA methylome, histone modification profiles as well as chromatin accessibility of SSCs have been revealed (Chen et al., 2020; Cheng et al., 2020; Guo et al., 2017; Hammoud et al., 2014; Jan et al., 2017; Lesch, Silber, McCarrey, & Page, 2016; Maezawa, Yukawa, Alavattam, Barski, & Namekawa, 2018; Sharma, Wistuba, Pock, Schlatt, & Neuhaus, 2019). Despite all these, the molecular mechanisms for SSC maintenance and development remain incompletely understood.

SSCs are able to undergo both self-renewal and differentiation. Unlike the self-renewal that replenishes the SSC and progenitor pool, the differentiation is an irreversible process committed to meiosis, which is stringently modulated by the stages of the seminiferous epithelium in the testis (De Rooij & Griswold, 2012; de Rooij & Russell, 2000). Typically, when they start to differentiate, SSCs and progenitors are gradually preparing their genome to later undergo a series of events in meiosis, such as initiation of double-strand breaks (DSBs), alignment, pairing and synapsis of homologous chromosomes, homologous recombination and formation of crossovers (Jan et al., 2012). It has thus been assumed that the preparations for these meiotic events in differentiating spermatogonia (Di-SG) are accompanied by dramatic alterations in chromatin structure. Traditional histological studies have revealed that Di-SG are equipped with increasing amount of condensed chromatin, namely heterochromatin, that rims the nucleus (Chiarini-Garcia & Russell, 2001, 2002). Despite this, the three-dimensional (3D) chromatin architecture of SSCs and Di-SG, and the higher-order chromatin dynamics during spermatogonial differentiation, have not been systematically studied.

The recently developed high-throughput chromosome conformation capture (Hi-C) technique enables detection and visualization of the dynamic chromatin, providing a desirable means to study the higher-order chromatin architecture and the key principles of genome packaging at the molecular level (Belton et al., 2012; Lieberman-Aiden et al., 2009). The higher-order chromatin can be spatially packaged into a hierarchy of the 3D genome, including A/B compartments, topologically associating domains (TADs) and chromatin loops (Dixon et al., 2012a; Rao et al., 2014), further influencing numerous DNA-related biological processes such as transcription, DNA replication and repair, mitotic and meiotic cell cycle progress, etc. (Gorkin, Leung, & Ren, 2014; Smallwood & Ren, 2013). Here, by using *in situ* Hi-C, RNA-seq and chromatin immunoprecipitation-sequencing (ChIP-seq), we systematically investigated the 3D chromatin architecture of Un-SG (which consist of SSCs and progenitors) and Di-SG, with an aim to unravel the higher-order chromatin dynamics during spermatogonial differentiation, as well as the regulation in gene transcription. We performed the studies on pigs, since pigs are an increasingly prevalent animal model in fundamental and translational research due to the resemblance to humans concerning anatomy, physiology, genetics and reproductive maturation (Swindle, Makin, Herron, Clubb, & Frazier, 2012; Voigt et al., 2021). Moreover, it is feasible to obtain a vast number of spermatogonial subpopulations from porcine testes for subsequent advanced bioinformatic analyses. We gained novel insights into the changing chromatin dynamics during spermatogonial differentiation that have so far not been reported. For instance, we identified that diminished higher-order chromatin architecture in meiotic cells, as shown by recent reports, is actually preprogramed in Di-SG. Besides, we unraveled the contribution of promoter-enhancer interactions (PEIs) to pre-meiotic transcriptional regulation, which has not been accomplished in previous studies due to limited cell input and resolution. Together, our study uncovered the 3D chromatin structure of SSCs/progenitors and Di-SG, as well as the interplay between higher-order chromatin architecture and dynamic gene expression during spermatogonial differentiation, which is expected to better the biological understanding of SSC self-renewal and differentiation and have implications for diagnosis and treatment of male sub-/infertility.

## Results

### Dynamic 3D chromatin architecture during spermatogonial differentiation

To uncover the 3D chromatin structure of SSCs/progenitors and Di-SG and the higher-order chromatin dynamics during spermatogonial differentiation, we first collected Un-SG and Di-SG from porcine testes. Our recent study has shown that SSEA4 is a surface marker of porcine Un-SG and that it can be used to enrich porcine Un-SG including transplantable SSCs with unprecedented efficiency (Zhang et al., 2020). Hence, Un-SG were isolated from 90-day-old porcine testes and enriched by fluorescence-activated cell sorting (FACS) employing an antibody against SSEA4, while Di-SG were isolated from 150-day-old porcine testes and enriched with a velocity sedimentation approach (STA-PUT) (Bryant, Meyer-Ficca, Dang, Berger, & Meyer, 2013; Liu et al., 2015). The high purity of collected spermatogonial subpopulations was validated by immunofluorescence staining and subsequent quantification of cells positive for stage-specific markers (Fig 1A). Then, we performed *in situ* Hi-C and RNA-seq analyses on the collected spermatogonial samples. For Hi-C analysis we generated high quality datasets from sufficient biological samples (8 independent samples for Un-SG and 8 for Di-SG), and obtained around 6.3 billion valid interactions for the overall 16 samples, with an average of 392 million valid interactions per sample (Table S1). For RNA-seq analysis we constructed transcriptomic libraries from 6 samples (3 independent samples for each spermatogonial subtype), with approximately 76 million paired reads per sample (Table S1).

**Fig 1.**
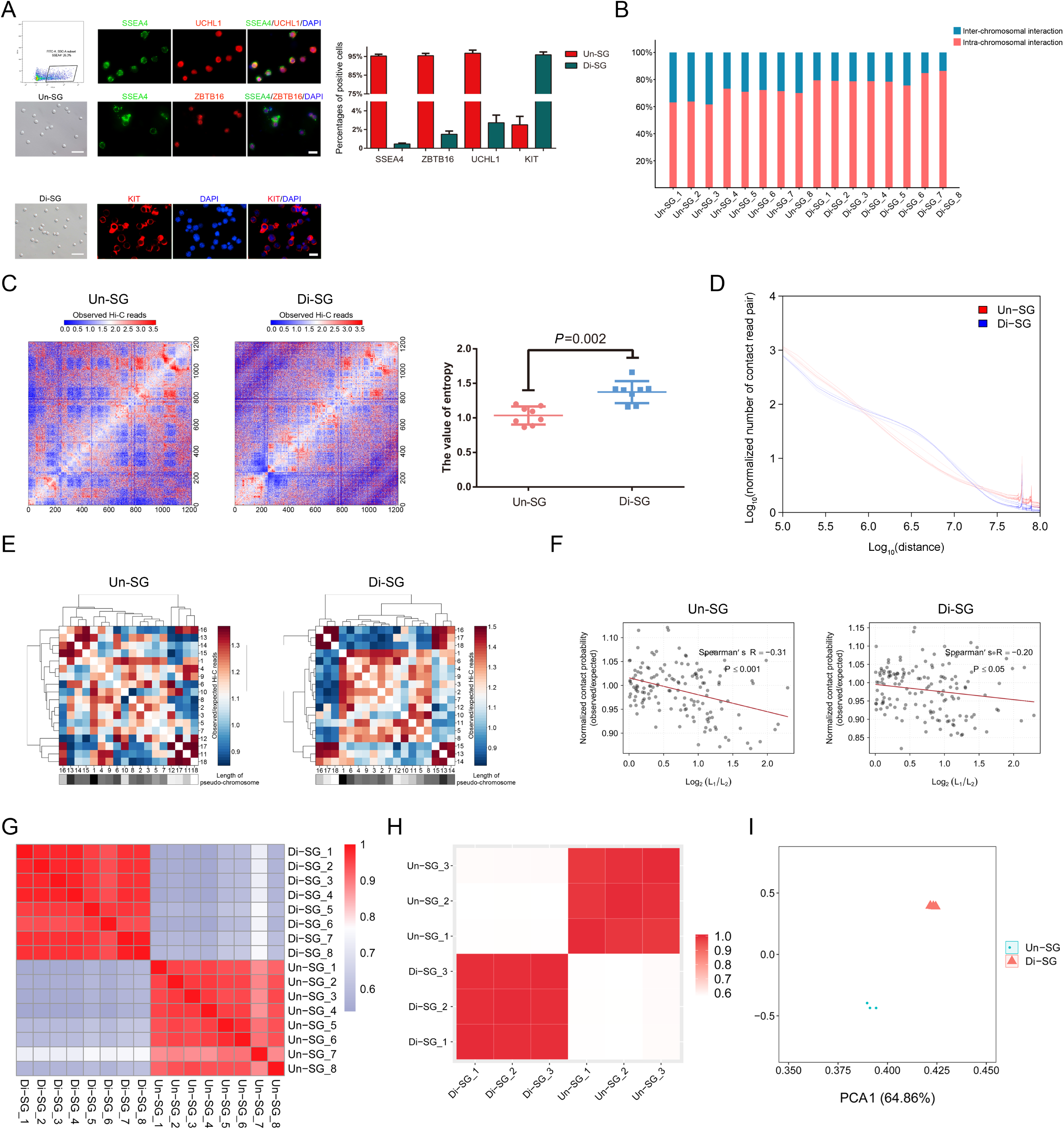
Dynamic 3D chromatin architecture during spermatogonial differentiation. (A) Enrichment and characterization of spermatogonial subpopulations. Un-SG were enriched by FACS employing an antibody against SSEA4, and both cell populations were subjected to immunofluorescence staining and quantification of cells positive for stage-specific markers (SSEA4, ZBTB16 and UCHL1 for Un-SG and KIT for Di-SG). Bar=50µm (brightfield) or 10µm (immunofluorescence). Data are presented as the mean ± SEM of eight independent samples. (B) The inter- and intra-chromosomal interaction ratios in all Un-SG and Di-SG samples. (C) The entropy difference between Un-SG and Di-SG. The intra-chromosome log_2_ Hi-C matrices are shown at 100kb resolution for chromosome 7. Data are presented as the mean ± SD of eight independent samples. *P*: Mann-Whitney U test, one-tailed. (D) The *P*(s) curves of Un-SG and Di-SG showing the interaction probability patterns between bin pairs at defined genomic distances. (E) The observed/expected number of contacts between any pair of 18 autosomes. The plaids with differential gray scale indicate the length of each chromosome. (F) The observed/expected number of interactions between any pair of 18 autosomes plotted against the length difference of these chromosomes. L_1_ or L_2_ refers to the length of chromosome (L_1_>L_2_), and length difference is indicated by log_2_(L_1_/L_2_). The dotted line represents the linear trend for obtained value. (G) HiC-Rep analysis illustrating the correlation of normalized Hi-C interaction matrices between Un-SG and Di-SG samples. (H) Pearson correlation analysis illustrating the correlation of transcriptomic data between Un-SG and Di-SG samples. (I) PCA plot showing the transcriptomic profiles of Un-SG and Di-SG samples.

The chromosomal conformation profiles revealed that compared with Un-SG, Di-SG exhibited increased intra-chromosomal interaction ratio (68.3% in Un-SG vs 80.2% in Di-SG) but decreased inter-chromosomal interaction ratio (31.7% in Un-SG vs 19.8% in Di-SG, Fig 1B, Table S1). When applying entropy as a measurement of the order in chromatin configuration (Seaman & Rajapakse, 2018), we observed that Di-SG had higher entropy (Fig 1C), suggesting more disorder in Di-SG. Next, we conducted a *P*(s) analysis to gain the interaction probability patterns between bin pairs at defined genomic distances (Naumova et al., 2013), and identified that Di-SG exhibited higher interaction probabilities than Un-SG at the distances between 1 and 10Mb, but that the trend was reversed at long distances (Fig 1D), in line with previous reports in mice and rhesus monkeys that the more advanced pachytene spermatocytes also displayed stronger interactions at short distances (between 1 and 5Mb) but weaker interactions at long distances (>10Mb) than spermatogonia (Luo et al., 2020; Wang et al., 2019).

We further analyzed the inter-chromosomal interaction. As expected, Un-SG and Di-SG exhibited similar nonrandomly distributed chromosomal positions. Longer chromosomes preferentially interacted with each other, and the same for shorter ones (Fig 1E). The negative correlation between the inter-chromosomal interaction probability and the chromosomal length also applied to both spermatogonial populations (Fig 1F), in accordance with recent findings in adipocytes and myoblasts (He et al., 2018).

Subsequently, we studied the normalized Hi-C interaction matrices with 100kb bin size for all independent samples. By employing HiC-Rep we detected a substantially lower correlation coefficient between Un-SG and Di-SG (*r*=0.60), in contrast with high correlation coefficients between ingroup samples (*r*=0.93 for Un-SG and *r*=0.96 for Di-SG, Fig 1G). These patterns can be validated by using QuASAR-Rep (Fig S1A), GenomeDISCO (Fig S1B), the Pearson correlation (Fig S1C) or principal component analysis (PCA, Fig S1D), corroborating distinct 3D chromatin organizations in two spermatogonial subgroups. In addition, we observed difference in transcriptomes between Un-SG and Di-SG, as reflected by a low correlation between Un-SG and Di-SG (*r*=0.58), in spite of the high correlation between ingroup samples (*r*=0.988 for Un-SG and *r*=0.997 for Di-SG, Fig 1H and 1I). Hence, these data indicate that alterations of chromatin configuration are accompanied by transcriptomic variations during spermatogonial differentiation.

### A/B compartment switches and changes during spermatogonial differentiation

Higher-order chromatin can be divided into A and B compartments, representing open chromatin regions with active genes and closed chromatin regions with inactive genes, respectively (Lieberman-Aiden et al., 2009). We then explored the A/B compartment index (20kb bin size) in autosomes of Un-SG and Di-SG. Pearson correlation analysis illustrated a low correlation of global A/B compartment states between Un-SG and Di-SG (*r*=0.78), in contrast with high correlations between ingroup samples (*r*=0.96 for Un-SG and *r*=0.95 for Di-SG, Fig 2A), which was validated by PCA analysis (Fig 2B). Despite the similar A/B compartment organization between Un-SG and Di-SG (Fig 2C), the compartment strength was decreased in Di-SG (Fig 2D and 2E), indicative of weakened compartmentalization and more disorder in Di-SG.

**Fig 2.**
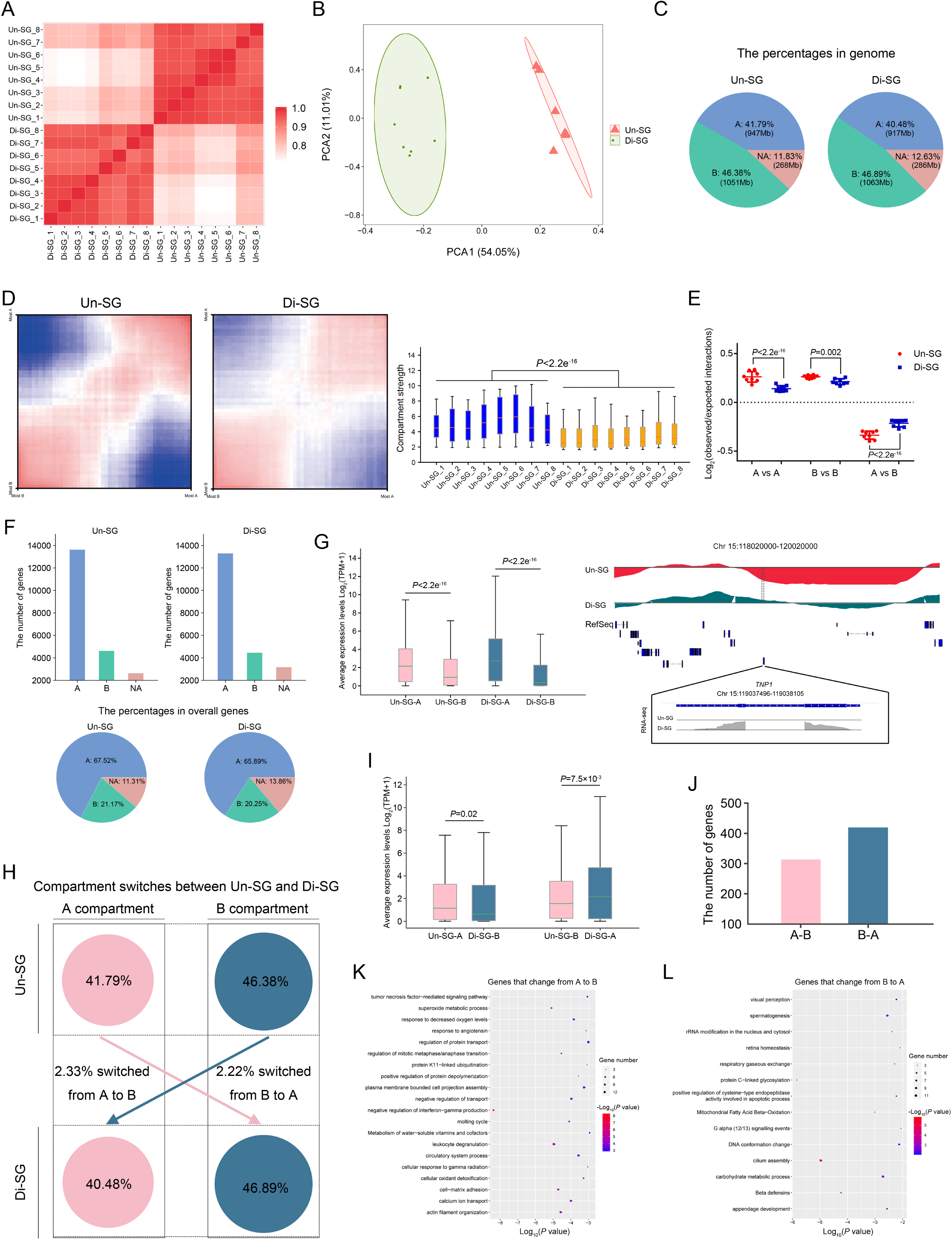
A/B compartment switches during spermatogonial differentiation. (A) Pearson correlation analysis illustrating the correlation of A/B indices between Un-SG and Di-SG samples. (B) PCA plot showing the A/B index profiles of Un-SG and Di-SG samples. (C) The proportions and lengths of A/B compartments in genome. (D) Left: saddle plot showing the compartment strength in chromosome 9. Right: the compartment strength in all Un-SG and Di-SG samples, defined as the A-A and B-B compartment interaction strength relative to the A-B compartment interaction strength. *P*: Mann-Whitney U test, one-tailed. (E) The interaction strength between A-A, B-B or A-B compartments. Data are presented as the mean ± SD of eight independent samples. *P*: Mann-Whitney U test, one-tailed. (F) The numbers (upper panel) and proportions (lower panel) of genes in A/B compartments. (G) Left: the average expression levels of genes in A/B compartments in Un-SG or Di-SG. *P*: Mann-Whitney U test, one-tailed. Right: PCA1 (the first eigenvalues, the upper part) and RefSeq view (the middle part) of chromosome 15, 118020000-120020000, as well as RNA-seq coverage track of *TNP1* (chromosome 15, 119037496-119038105, the lower part) showing that *TNP1*, which was upregulated during spermatogonial differentiation, was located in the B compartment in Un-SG but switched to the A compartment in Di-SG. PCA1 was calculated via eigenvector decomposition on the observed/expected intra-chromosomal interaction matrices. (H) A schematic overview illustrating the proportions of genomic regions subjected to A/B compartment switches (A to B or B to A) between Un-SG and Di-SG. (I) The average expression levels of genes that changed from A to B or from B to A. *P*: Mann-Whitney U test, one-tailed. (J) The numbers of genes that changed from A to B or from B to A. (K and L) Gene ontology-biological process (GO-BP) analysis of genes that changed from A to B (K) or from B to A (L).

In both cell populations, the A compartments harbored the majority of genes (Fig 2F), and genes in the A compartments showed higher expression levels than those in the B compartments (Fig 2G). Previous studies have reported the correlation of the A/B compartment switch with transcriptional regulation (Lieberman-Aiden et al., 2009). We thus probed the occurrence of the A/B compartment switch during spermatogonial differentiation. We found that 52.88Mb and 50.22Mb, making up 2.33% and 2.22% of the autosomal genome, underwent the A-B and B-A switch, respectively (Fig 2H). Genes that change from compartment A to B during spermatogonial differentiation tended to show lower expression levels in Di-SG than in Un-SG, whereas genes that change from B to A tended to be upregulated (Fig 2I). Specifically, 314 genes that change from compartment A to B (Fig 2J, Table S2) were enriched in cell-matrix adhesion, regulation of mitotic metaphase/anaphase transition and response to decreased oxygen levels (Fig 2K, Table S3), whereas 420 genes that change from B to A (Fig 2J, Table S2) fell in terms such as carbohydrate metabolic process, spermatogenesis and DNA conformation change (Fig 2L, Table S3). For instance, *ATM*, a protein involved in DNA damage response and located in the A compartment in Un-SG, was downregulated (FDR<0.05, fold change>2) in Di-SG where it was located in the B compartment. *TNP1*, which plays important roles in spermiogenesis and was significantly upregulated during spermatogonial differentiation, was located in the B compartment in Un-SG but switched to the A compartment in Di-SG (Fig 2G, Table S2).

Compartments experiencing the A-A or B-B change can refer to those correlated with increasingly open or closed chromatin, respectively. We found that 44.2Mb and 65.2Mb, accounting for 1.95% and 2.88% of the autosomal genome, were subjected to the A-A and B-B change, respectively (Fig S2A). Genes that undergo the compartment A-A change during spermatogonial differentiation tended to show higher expression levels in Di-SG than in Un-SG, whereas genes that undergo the B-B change tended to be downregulated (Fig S2B). Specifically, 739 genes that undergo the compartment A-A change (Fig S2C, Table S2) were enriched in male gamete generation, regulation of mitotic/meiotic cell cycle and chromosome organization (Fig S2D, Table S3), whereas 244 genes that undergo the B-B change (Fig S2C, Table S2) fell in terms such as cellular response to hormone stimulus, regulation of cell adhesion and Notch signaling pathway (Fig S2E, Table S3). Genes that undergo the compartment A-A change included *HSPA2*, *STRA8*, *SOX30*, *MLH1* and *METTL3*, all of which have been reported to be involved in spermatogenesis and showed higher expression levels in Di-SG than in Un-SG (FDR<0.05, fold change>2). *YTHDC2*, an N^6^-methyladenosine (m^6^A)-binding protein playing regulatory roles in spermatogenesis, underwent the B-B change and was downregulated (FDR<0.05, fold change>2) during spermatogonial differentiation (Table S2). Together, our data suggest that switches and changes of A/B compartments play important regulatory roles in dynamic gene expression during spermatogonial differentiation.

### TAD dynamics during spermatogonial differentiation

TADs have been reported to be generally conserved among distinct cell types (Fraser et al., 2015; He et al., 2018; Nora et al., 2012; Rubin et al., 2017; Siersbaek et al., 2017). Nevertheless, recent articles reported reorganization of TADs during spermatogenesis (Alavattam et al., 2019; Luo et al., 2020; Patel et al., 2019; Vara et al., 2019; Wang et al., 2019). Drastic alterations of TADs have also been identified in oogenesis, i.e. TADs undergo gradual attenuation and then vanish during oogenesis, and it is only until the two-cell or even later embryo developmental stage that TADs reemerge and gradually restore (Ke et al., 2017), suggesting distinctive TAD dynamics in gametogenesis and early embryo development. Yet, whether TADs are conserved or subjected to alterations during spermatogonial differentiation remains to be explored. To this end, we analyzed the TAD architecture at 20kb resolution in both spermatogonial subtypes. We identified that TADs constituted the majority of the genome (Fig 3A), with a decrease of the TAD number but an increase of the mean TAD size (Fig 3B) during spermatogonial differentiation. We observed a low correlation between Un-SG and Di-SG (0.80), in contrast with high correlations of TAD architecture between ingroup samples (0.91 for Un-SG and 0.86 for Di-SG), as reflected by Jaccard indices (Fig 3C). Then, we employed the insulation score (IS, Fig 3D), the directional index (DI, Fig 3E) as well as aggregate Hi-C maps (Fig 3F) to measure the strengths of TAD boundaries, and found all of them declined in Di-SG. Moreover, the domain score (D-score), which is defined by the ratio of intra-TAD interactions in the overall intra-chromosomal interactions (Krijger et al., 2016) and able to quantify the tendency of TADs to self-interact (Stadhouders et al., 2018), was decreased in Di-SG (Fig 3G), further suggesting that TADs are weakened during spermatogonial differentiation. Thus, our data complement previous studies by showing that TAD attenuation already initiates at the pre-meiotic spermatogonial differentiation stage, towards the TAD dissolution occurring in subsequent meiosis (Alavattam et al., 2019; Luo et al., 2020; Patel et al., 2019; Vara et al., 2019; Wang et al., 2019).

**Fig 3.**
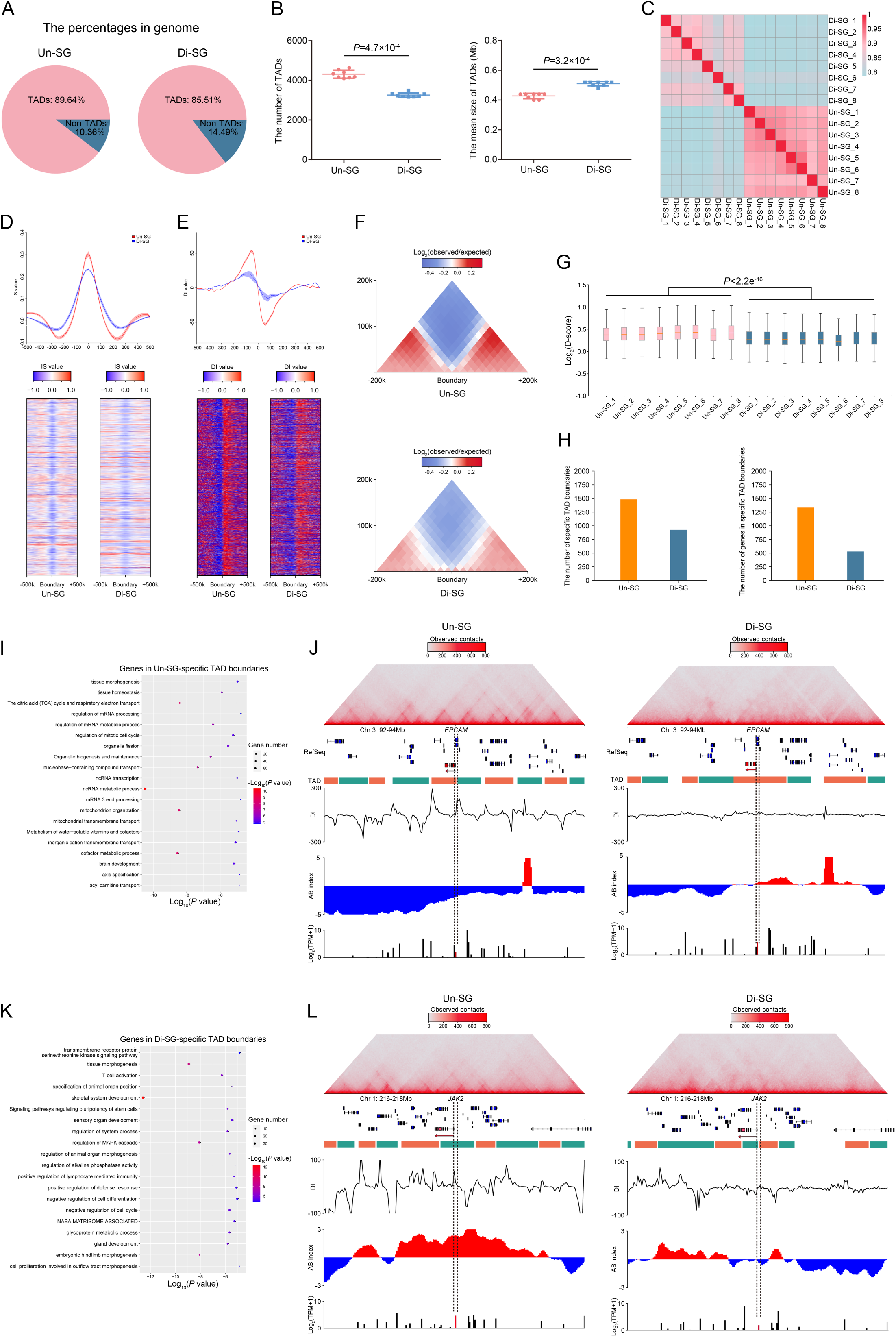
TAD dynamics during spermatogonial differentiation. (A) The proportions of TADs and non-TADs in genome. (B) The numbers (left) and mean sizes (right) of TADs in Un-SG and Di-SG. Data are presented as the mean ± SD of eight independent samples. *P*: Mann-Whitney U test, one-tailed. (C) Jaccard indices illustrating the correlation of TAD architecture between Un-SG and Di-SG samples. (D and E) The mean IS (D) and DI (E) value of TADs and the flanking regions (±500k) in Un-SG and Di-SG. (F) The aggregate Hi-C map showing the average observed/expected chromatin interaction frequencies at TADs and the flanking regions (±200k) in Un-SG and Di-SG. (G) The D-score in all Un-SG and Di-SG samples. *P*: Mann-Whitney U test, one-tailed. (H) The numbers of specific TAD boundaries (left) and their harbored genes (right) in Un-SG and Di-SG. (I) GO-BP analysis of genes in Un-SG-specific TAD boundaries. (J) Views of the observed/expected chromatin interaction frequencies (the upper panel), RefSeq (the middle panel), DI, A/B index and RNA-seq coverage (the lower panel) at chromosome 3, 92-94Mb, revealing that *EPCAM* was harbored in Un-SG-specific TAD boundaries, in B-A switching compartments and upregulated in Di-SG. (K) GO-BP analysis of genes in Di-SG-specific TAD boundaries. (L) Views of the observed/expected chromatin interaction frequencies (the upper panel), RefSeq (the middle panel), DI, A/B index and RNA-seq coverage (the lower panel) at chromosome 1, 216-218Mb, revealing that *JAK2* was harbored in Di-SG-specific TAD boundaries, in A-B switching compartments and downregulated in Di-SG.

Later, we investigated whether TAD attenuation contributes to dynamic gene expression during spermatogonial differentiation. There were 1482 Un-SG-specific TAD boundaries harboring 1333 genes (Fig 3H, Table S4) that are related to mitochondrion organization, regulation of mRNA metabolic process and mitotic cell cycle (Fig 3I, Table S5). These genes included spermatogonial markers (e.g., *TSPAN33*, *EPCAM*, Fig 3J), those involved in spermatogonial self-renewal (e.g., *FOXO1*), in differentiation (e.g., *BMP4*, *DAZL*, *WNT3A*) and in meiosis (e.g., *SPO11*, Table S4). By contrast, 529 genes, which were embedded in 926 Di-SG-specific TAD boundaries (Fig 3H, Table S4), were implicated in important biological processes during spermatogonial development, such as regulation of MAPK cascade, transmembrane receptor protein serine/threonine kinase signaling pathway, and regulation of cell cycle and cell differentiation (Fig 3K, Table S5). Genes falling in this group included *ZBTB16*, a pivotal transcriptional regulator of spermatogonial self-renewal, those involved in pluripotency maintenance (e.g., *LIF*), and the JAK/STAT signaling component *JAK2* (Fig 3L, Table S4).

Long-range interacting TADs have been reported to be able to form TAD interaction networks, namely TAD cliques, to influence lineage-specific differentiation (Paulsen et al., 2019). It would be intriguing to explore whether the TAD interaction network, other than the TAD itself, is also attenuated during spermatogonial differentiation. Hence, we defined a TAD clique as a cluster of five or more interacting TADs in our Hi-C data. We observed high correlations of TAD cliques between ingroup samples (Pearson’s *r*=0.77 for Un-SG and *r*=0.78 for Di-SG, Fig S3A). In accordance with the variation of TAD boundaries, the formation of TAD cliques (reflected by the number of TAD cliques, Fig S3B), the genome coverage by TAD cliques (Fig S3C) as well as the percentage of TADs in cliques (Fig S3D) were diminished in Di-SG, indicating attenuation of the TAD interaction network during spermatogonial differentiation. Besides, we found that TAD cliques were enriched in B compartments relative to A compartments in both spermatogonial populations (Fig S3E). Thus, the attenuation of TAD cliques in Di-SG suggests the facilitated transcription during spermatogonial differentiation.

### Identification of PEIs and their regulation in gene expression during spermatogonial differentiation

Chromatin can be spatially packaged into the 3D genome architecture chromatin loops, facilitating the interactions between promoters and distant DNA regulatory elements. In this way, long-range enhancers are able to physically contact with the target promoters, modulating the temporal and spatial expression of target genes (Mifsud et al., 2015; Schoenfelder et al., 2015). To gain knowledge about the potential PEIs and their regulation in dynamic gene expression during spermatogonial differentiation, we combined the reads from 8 independent samples of Un-SG or Di-SG into single sets of Hi-C data (to reach the resolution of 5kb). We identified overall 67064 PEIs in Un-SG and 60344 in Di-SG (Fig 4A), and that the 15679 and 14032 promoters interacted with at least one enhancer in Un-SG and Di-SG, respectively (Fig 4B). The majority of PEIs were within 100kb (Fig 4C), implicated skipping enhancers (enhancers that are not the closest to promoters, Fig 4D) and, as expected, within TADs (Fig 4E) in both spermatogonial subtypes. Besides, we detected slightly more enhancers that interact with each promoter in Un-SG than in Di-SG (Fig 4F).

**Fig 4.**
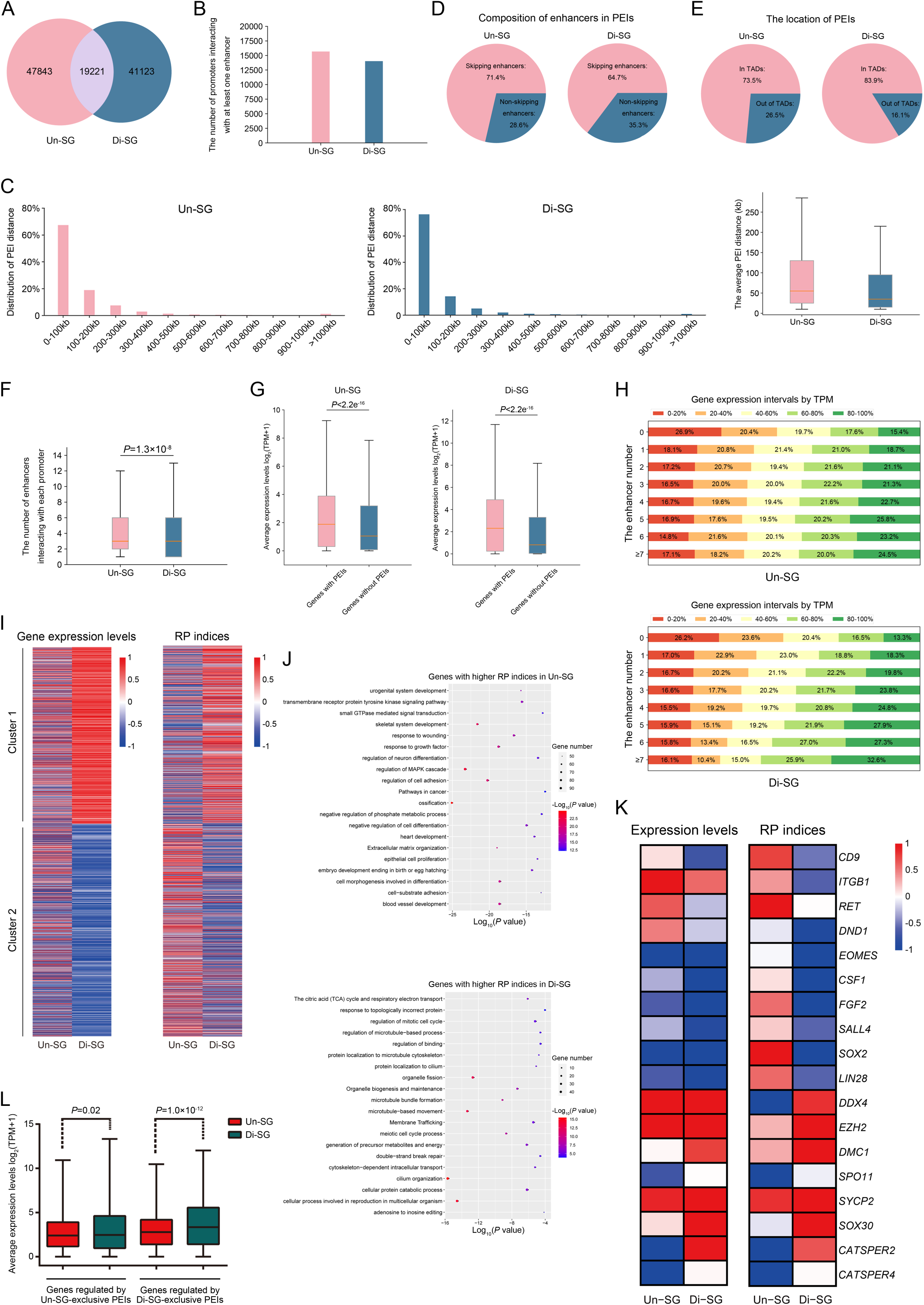
PEI regulation in gene expression during spermatogonial differentiation. (A) The numbers of PEIs in Un-SG and Di-SG samples. The number in the overlapped region refers to PEIs present in both populations. (B) The numbers of promoters that interact with at least one enhancer in Un-SG and Di-SG samples. (C) Left: distribution of PEI distance in Un-SG and Di-SG samples. Right: the average PEI distance in Un-SG and Di-SG samples. (D) Composition of skipping and non-skipping enhancers in PEIs. (E) The proportions of PEIs in or out of TADs. (F) The average numbers of enhancers that interact with each promoter in Un-SG and Di-SG samples. *P*: Mann-Whitney U test, one-tailed. (G) The average expression levels of genes with or without PEIs. *P*: Mann-Whitney U test, one-tailed. (H) More interacting enhancers are associated with higher proportions of genes in top gene expression intervals. The numbers in columns refer to the proportions of genes in each gene expression interval. (I) Heatmaps of expression levels and RP indices of two clusters of genes that were expressed in either or both spermatogonial subpopulations (TPM>1) and that exhibited a fold change of ≥4. (J) GO-BP analysis of genes with differential RP indices during spermatogonial differentiation. (K) Heatmaps of representative genes with differential RP indices and expression levels during spermatogonial differentiation. (L) The average expression levels of genes regulated by Un-SG- or Di-SG-exclusive PEIs. *P*: Mann-Whitney U test, one-tailed.

As expected, genes with PEIs exhibited generally higher expression levels than those without PEIs (Fig 4G), and more interacting enhancers were also associated with higher gene expression in both Un-SG and Di-SG (Fig 4H), suggesting cumulative effects of enhancers on the transcriptional levels of target genes. Since the multiple enhancers that interact with promoters vary in interacting intensity, we then introduced a regulatory potential (RP) index that combines both the number and intensity of the interacting enhancers to quantify their potential for transcriptional regulation of target genes. We identified that alterations of the RP index are generally consistent with gene transcriptional changes (Fig 4I), suggesting that PEIs orchestrate transcription during spermatogonial differentiation. Later, we detected that the 3985 genes showed significantly higher RP indices in Un-SG than in Di-SG (RP_Un-SG_-RP_Di-SG_>3, fold change>2, Table S6). These genes were related to regulation of cell adhesion, response to growth factor and cell morphogenesis involved in differentiation (Fig 4J, Table S7). Examples in this case are undifferentiated spermatogonial markers (*CD9*, *ITGB1*), genes important to spermatogonial self-renewal (*RET*, *DND1*, *EOMES*, *CSF1*, *FGF2*, *SALL4*) and pluripotency (*SOX2*, *LIN28*, Fig 4K, Table S6) that exhibited both higher RP indices and expression levels (FDR<0.05, fold change>2) in Un-SG than in Di-SG. By contrast, 2741 genes, falling in terms such as cellular process involved in reproduction in multicellular organism, meiotic cell cycle process, regulation of mitotic cell cycle and DSB repair (Fig 4J, Table S7), showed significantly lower RP indices in Un-SG than in Di-SG (RP_Un-SG_-RP_Di-SG_<-3, fold change>2, Table S6). Illustrations of this point are the pan-germ cell marker *DDX4*, genes involved in spermatogonial differentiation (*EZH2*), in meiosis (*DMC1*, *SPO11*, *SYCP2*), in spermiogenesis (*SOX30*) and in sperm motility (*CATSPER2*, *CATSPER4*, Fig 4K, Table S6) that displayed both lower RP indices and expression levels in Un-SG than in Di-SG, lending further support to the positive correlation between the RP index and gene expression.

In addition, we identified that some genes were regulated by spermatogonial subtype-exclusive PEIs, i.e., PEIs present in only Un-SG or Di-SG. To be more precise, 2730 genes, including those involved in spermatogenesis, such as *BMP4*, *DMRT1*, *RXRG*, *SYCP1*, *PRM1*, *PRM2*, *METTL3*, *BRCA2*, *WNT3A*, *FTO*, *NEDD4* and *GDNF* (Table S8), were regulated by Un-SG-exclusive PEIs, whilst 1671 genes, such as *CXCR4*, *LHX1*, *LIF*, *RARA*, *RARB*, *SPO11*, *ATR*, *SMC4*, *TNP2*, *PRM3* and *CATSPER4* (Table S8) that are also spermatogenesis-related, were with Di-SG-exclusive PEI regulation. As expected, genes regulated by cell type-exclusive PEIs were associated with elevated expression levels (Fig 4L). Taken together, these data indicate that PEIs act as an important element of transcriptional regulation during spermatogonial differentiation.

### Characterization of H3K27ac-marked active enhancers and their regulation in gene expression during spermatogonial differentiation

To delve into the role for enhancers in PEIs and further in gene expression, we performed ChIP-seq analyses on the collected spermatogonial samples, employing antibodies against H3K27ac (an active enhancer marker) or H3K4me3 (an active promoter marker). By combining Hi-C chromatin interactions and H3K27ac ChIP-seq data and applying the ROSE algorithms (Whyte et al., 2013), we defined regular enhancers (REs) and super enhancers (SEs), and by splitting each SE into 5kb bins, we further classified SEs into hierarchical (which are composed of hub and non-hub enhancers) and non-hierarchical enhancers, as previously reported (Huang et al., 2018). Arguably, substantially more REs than SEs were identified in both spermatogonial populations (Fig 5A). Then, we looked into the enhancers within PEIs, and found that in both subgroups only a small fraction of enhancers fell in the scope of SEs (Fig 5B). Various enhancers tended to interact with active promoters (marked by H3K4me3, Fig 5C). Consistently, when looking into genes with PEIs, it could be found that only a minor fraction of genes was regulated by SEs (Fig 5D), even though most genes had active promoters interacting with enhancers (Fig 5E).

**Fig 5.**
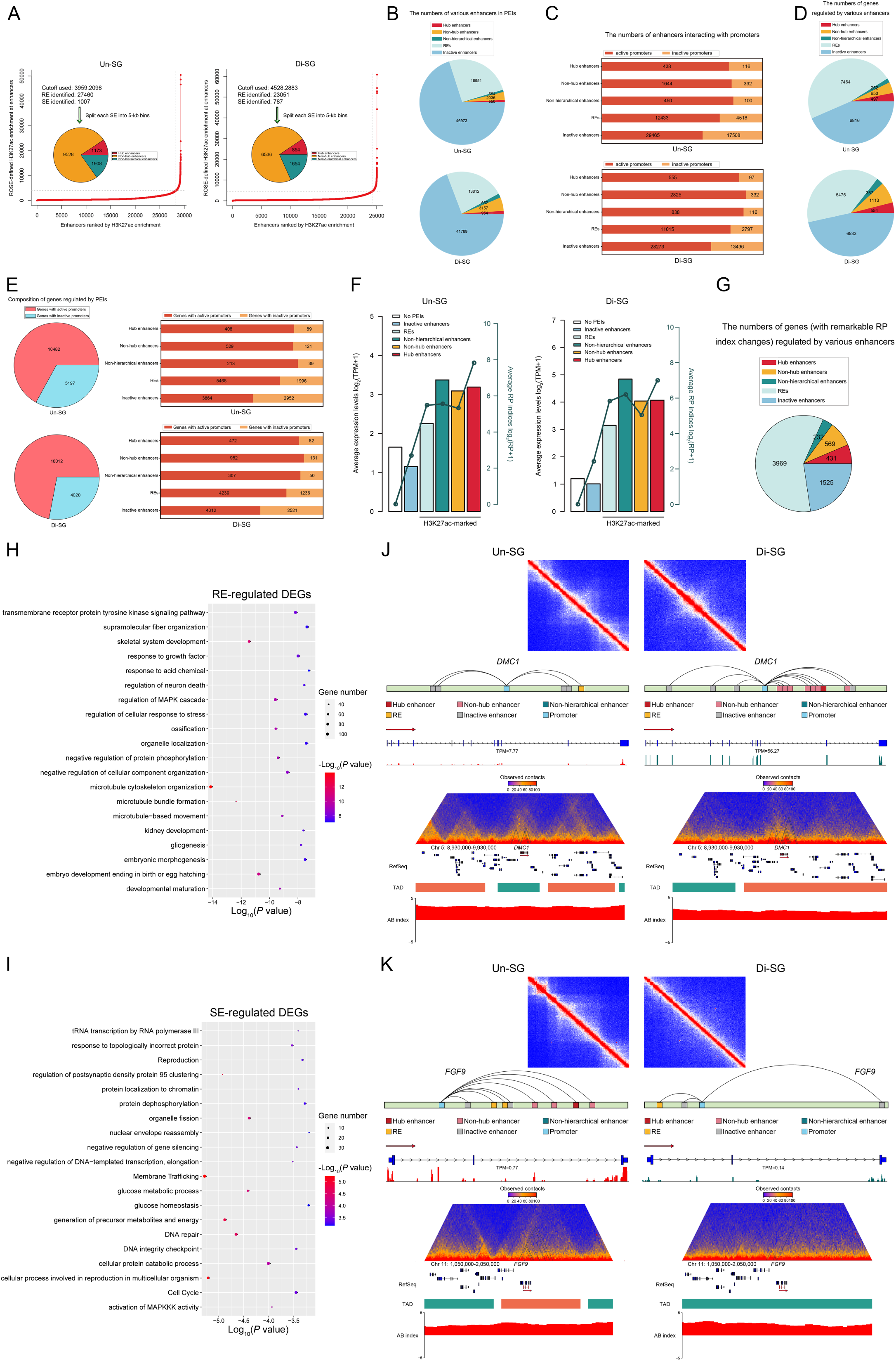
Regulation of H3K27ac-marked active enhancers in gene expression during spermatogonial differentiation. (A) Definition and numbers of REs and hierarchically organized SEs in two spermatogonial populations. (B) The numbers of different categories of enhancers within PEIs. (C) The numbers of different categories of enhancers that interact with active or inactive promotors. (D) The numbers of PEI genes regulated by different categories of enhancers. (E) Left: the numbers of PEI genes with active or inactive promoters. Right: the numbers of PEI genes regulated by various enhancers and promoters. (F) Average expression levels and RP indices (indicated by bars and lines, respectively) of genes regulated by different categories of enhancers. (G) The numbers of genes (with remarkable RP index changes) regulated by different categories of enhancers. (H and I) GO-BP analysis of RE (H) and SE (I)-regulated genes with differential RP indices and expression levels during spermatogonial differentiation. (J and K) The upper panel: the contact matrix showing stripes at the *DMC1* (J) and *FGF9* (K) SE loci. The middle panel: models for PEI regulation of *DMC1* (J) and *FGF9* (K), as well as RNA-seq coverage at their genomic loci. The lower panel: views of the observed/expected chromatin interaction frequencies, RefSeq, TAD and A/B index at chromosome 5: 8,930,000-9,930,000 (J) and chromosome 11: 1,050,000-2,050,000 (K).

It has been reported that various enhancers differentially contribute to gene expression (Hnisz et al., 2013; Huang et al., 2018). As expected, we found that in both spermatogonial populations genes with higher RP indices and expression levels tended to be regulated by active enhancers that are marked by H3K27ac (i.e., REs and SEs), and that genes regulated by SEs showed generally higher expression levels than those targeted by REs (Fig 5F). RE and SE-associated genes were also prone to have active promoters (marked by H3K4me3, Fig 5E, right), together acting as hallmarks of active transcription. Further analysis revealed that of the genes showing remarkable RP index changes during spermatogonial differentiation (RP_Un-SG_-RP_Di-SG_>3 or <-3, fold change>2), 59.0% (3969/6726) were regulated by REs (Fig 5G), of which 54.3% (2154/3969) showed differential expression levels (FDR<0.05, fold change>2) in two spermatogonial populations (Table S9). These genes were enriched with regulation of MAPK cascade, transmembrane receptor protein tyrosine kinase signaling pathway, response to growth factor and developmental maturation (Fig 5H, Table S10). By contrast, 18.3% (1232/6726) of the genes with remarkable RP index changes were regulated by SEs (Fig 5G), of which 55.0% (677/1232) were differentially expressed (Table S9). These genes fell in terms such as cellular process involved in reproduction in multicellular organism, generation of precursor metabolites and energy, DNA repair and activation of MAPKKK activity (Fig 5I, Table S10). Hence, these data suggest that both REs and SEs are important regulatory elements underlying the dynamic transcriptome during spermatogonial differentiation.

Of the active enhancers that are marked by H3K27ac, SEs particularly appealed to us, due to their substantially high levels of activity, enrichment of active chromatin characteristics, as well as pivotal roles in cell fate determination (Hnisz et al., 2013; Parker et al., 2013; Whyte et al., 2013). To gain more insights into SE-mediated transcriptional regulation during spermatogonial differentiation, we inspected 1232 SE-associated genes with remarkable RP index changes. Of these, 35.0% (431/1232), 46.2% (569/1232) and 18.8% (232/1232) were regulated by hub enhancers, non-hub enhancers and non-hierarchical enhancers, respectively (Fig 5G, Table S9). Intriguingly, we found that hub enhancer-targeted genes harbored some with well-known roles in spermatogenesis. For instance, *DMC1*, a gene essential for DSB repair and meiotic homologous recombination (Kagawa & Kurumizaka, 2010), was targeted by hub enhancers only in Di-SG and also upregulated in Di-SG (FDR<0.05, fold change>2, Fig 5J, Table S9). Another illustration of this point is *CATSPER2*, a member of the *CATSPER* gene family with functions in sperm motility (Visser et al., 2011) (Fig S4A, Table S9). *FGF9* is a downstream gene of *SOX9* that plays crucial roles in testicular development. Recently, a novel role for *FGF9* in promoting SSC self-renewal has been reported (Yang et al., 2020). Interestingly, this gene was targeted by hub enhancers only in Un-SG and showed higher expression levels in Un-SG (Fig 5K, Table S9). Genes with hub enhancer regulation also included *DND1*, a gene encoding DND1 which associates with NANOS2 to promote spermatogonial self-renewal (Niimi et al., 2019) (Fig S4B, Table S9), as well as *EZH2*, an epigenetic factor capable of modulating spermatogonial differentiation and apoptosis (Jin et al., 2017) (Fig S4C, Table S9). These results thus corroborate that SEs, in particular hub enhancers, act as important regulatory elements of dynamic gene transcription during spermatogonial differentiation.

## Discussion

Several recent articles reported reorganized chromatin architecture during mammalian spermatogenesis. As reported, compartmentalization, TADs or loops underwent dissolution and reestablishment with spermatogenic cell development, and gene transcription seemed to be independent of the chromatin structure at certain stages such as the pachytene stage (Alavattam et al., 2019; Luo et al., 2020; Patel et al., 2019; Vara et al., 2019; Wang et al., 2019). Here, by integrating Hi-C, RNA-seq and ChIP-seq data, we delved into the higher-order chromatin structural dynamics and their influences upon transcriptional regulation during spermatogonial differentiation. Our findings complement previous studies by showing that the dynamic alterations in 3D chromatin organization already initiate at the pre-meiotic spermatogonial differentiation stage: Di-SG have increased disorder but weakened compartmentalization and TADs in comparison with Un-SG. Our results also suggest that A/B compartments and TADs are related to dynamic gene expression during spermatogonial differentiation. Moreover, since it is feasible to obtain a vast number of spermatogonial subpopulations from porcine testes, we proceeded to explore the contribution of PEIs to pre-meiotic transcriptional regulation, which has not been accomplished in previous studies due to limited cell input and resolution.

One of the most striking findings in this study might be the dynamic 3D chromatin structure during spermatogonial differentiation, which is in contrast with a recent article describing minimal alterations in higher-order chromatin architecture between primitive type A spermatogonia and type A spermatogonia in mice (Luo et al., 2020). From our perspective, the discrepancy could be ascribed to several respects. First, since peers and us have demonstrated that SSEA4 is a conserved surface marker of Un-SG and that it can be employed to enrich non-human primate, human and porcine Un-SG including transplantable SSCs efficiently (Fayomi & Orwig, 2018; Guo et al., 2017; Zhang et al., 2020), here we exploited an antibody against SSEA4 in conjunction with FACS to enrich Un-SG with high purity for subsequent bioinformatic analyses, distinct from Luo and colleagues who utilized a STA-PUT procedure to collect spermatogonial subpopulations with relatively lower purity (Luo et al., 2020). Second, as reported in that article, primitive type A spermatogonia were isolated from 6-day-old mice (Luo et al., 2020). Indeed, mouse male germ cells at this developmental stage consist of not only Un-SG but also a fraction of Di-SG likely committed to the first wave of spermatogenesis (Culty, 2013; Law & Oatley, 2020; Manku & Culty, 2015; Niedenberger, Busada, & Geyer, 2015), which are morphologically indistinguishable and not able to be separated by velocity sedimentation approaches such as STA-PUT, and due to this heterogeneity, the reported minimal alterations in higher-order chromatin architecture between the collected spermatogonial subgroups might be underrepresented for the probably changing chromatin dynamics during spermatogonial differentiation. Third, while it has traditionally been acknowledged that spermatogenesis is a generally conserved process among mammalian species, recent single-cell RNA-seq analyses of testicular cells from mice, human and non-human primates disclosed some divergent characteristics during mammalian spermatogenesis (Lau, Munusamy, Ng, & Sangrithi, 2020; Shami et al., 2020). Hence, the possibility remains that the 3D chromatin dynamics during spermatogonial differentiation *per se* differ between mice and pigs.

Previous studies have suggested that chromatin reorganization is a characteristic event during stem cell differentiation and lineage specification (Dixon et al., 2015; Ke et al., 2017; Paulsen et al., 2019). Here, we identified that also spermatogonial differentiation entailed dynamic alterations in 3D chromatin organization, characterized by increased disorder but attenuated compartmentalization and TADs. Spermatogonial differentiation has been known as a process that implicates pronounced transitions in cell-cycle, transcriptional and metabolic regulators, separating the largely quiescent SSCs (which principally rely on glycolysis for energy supply) from the more proliferative Di-SG (which preferentially utilize oxidative phosphorylation to produce ATP) (Caldeira-Brant et al., 2020; Chen et al., 2020; Guo et al., 2017; Lord & Nixon, 2020; Tan & Wilkinson, 2019, 2020). Weakened compartmentalization and more disorder in Di-SG might therefore be related to cell-cycle transitions and metabolic shifts. TADs have recently been reported to almost vanish in pachytene spermatocytes (Alavattam et al., 2019; Luo et al., 2020; Patel et al., 2019; Vara et al., 2019; Wang et al., 2019), even though extensive transcription occurs with dissolved TADs. Our data demonstrated that TAD attenuation already initiated at the pre-meiotic spermatogonial differentiation stage. This, along with the marked upregulation of many meiosis-related transcripts in Di-SG, as revealed by the present and previous studies (Chen et al., 2020; Jan et al., 2017; Zheng et al., 2018), corroborate that spermatogonial differentiation is for a large part a transitional process that gradually prepares the genome for the subsequent meiotic events.

Previous studies have also suggested the need to unravel the elusive and enigmatic relationship between transcription and chromatin configuration. Here, we identified that the dynamic gene expression during spermatogonial differentiation could be influenced by A/B compartment switches and changes, as well as TAD boundaries and cliques. To gain more knowledge about the contribution of delicate chromatin organization to gene transcription during this process, we probed PEIs in Un-SG and Di-SG under 5kb bins. We introduced a RP index to quantify the potential of interacting enhancers for transcriptional regulation. As expected, the RP index was found positively correlated with gene expression during spermatogonial differentiation. Our findings thus provide direct evidence that apart from epigenetic modification and non-coding RNAs, also PEIs are an important element of transcriptional regulation in the process of male germline development. Further, we characterized REs and SEs, and investigated the structural hierarchy of SEs on the basis of chromatin interactions in two spermatogonial populations. Intriguingly, we identified that several genes with well-known roles in spermatogenesis were potentially regulated by hub enhancers within hierarchical SEs, suggesting a role for the structural hierarchy of SEs in transcriptional regulation during spermatogonial differentiation. Future perturbation studies by using, e.g., the CRISPR-Cas9 strategy, will functionally validate the role of hub enhancers in SE structure and further in transcriptional regulation. Nevertheless, a prerequisite for this would be establishment of an optimized long-term culture system that enables stable propagation and induced differentiation of porcine SSCs, akin to their mouse counterparts that not only readily proliferate and differentiate *in vitro* but also seem amenable to CRISPR-Cas9-mediated genome editing (Sato et al., 2015; Wu et al., 2015; Zheng et al., 2017).

To sum up, we systematically investigated the 3D genome organization and its correlation with transcriptional regulation during spermatogonial differentiation. We identified that diminished higher-order chromatin architecture in meiotic cells, as shown by recent reports, is actually preprogramed in Di-SG, delineating unidirectional development of male germline, and have also for the first time, to our knowledge, unraveled the contribution of PEIs to pre-meiotic transcriptional regulation. Recent studies exploiting the single-cell RNA-seq technique uncovered the transcriptomes of progressive spermatogenic subtypes during mammalian spermatogenesis (Suzuki et al., 2019; Tan & Wilkinson, 2019, 2020). In future, development of single-cell Hi-C technology would help to unravel the finer 3D chromatin structural difference between SSCs and progenitors, enabling more comprehensive insights into the higher-order chromatin dynamics during male germline development, and with functional perturbation analyses, the roles of 3D genome conformation in transcriptional activity could be validated. Overall, the present study adds to the growing body of knowledge about chromatin configuration related to male fertility, and may potentially contribute to treatment of male infertility by SSC therapy, i.e., SSC auto-transplantation (Mulder et al., 2016) or *in vitro* differentiation into sperm (Lei et al., 2020).

## Materials and methods

### Testis samples

Testes were obtained from 90 or 150-day-old Duroc pigs (Besun farm, Yangling, Shaanxi, China). After surgical castration, testes were placed in Dulbecco’s phosphate-buffered saline (DPBS) supplemented with 2% penicillin/streptomycin (Hyclone) and transported to the laboratory on ice. All animal procedures were in accordance with and approved by the Institutional Animal Care and Use Committee of Northwest A&F University.

### Isolation and enrichment of spermatogonial populations

Un-SG were isolated from 90-day-old porcine testes and enriched by FACS employing an antibody against SSEA4. To obtain the single-cell suspension, the testis tunica albuginea and visible connective tissue were removed, and then exposed to Type IV Collagenase (2 mg/mL; Thermo Fisher Scientific) at 35°C for 20 minutes with periodic shaking. After three times of washing with DPBS to remove interstitial cells, the obtained seminiferous tubules were incubated with hemolytic fluid for 2 minutes to remove erythrocytes, followed by treatment with 0.25% trypsin-EDTA (Hyclone) at 37°C for 5 minutes to obtain the single-cell suspension. After centrifugation, the cell pellet was resuspended in Dulbecco’s modified Eagle medium (DMEM, high glucose; Hyclone) supplemented with 5% fetal bovine serum (FBS; Hyclone) and subjected to differential plating, as previously reported (Zhang et al., 2020). The suspension containing Un-SG was then collected and applied to FACS. In brief, the cells were washed with chilled FACS buffer (DPBS with 1% FBS and 2mM EDTA) and then incubated with the mouse anti-SSEA4 antibody (1: 50; 4755S, Cell Signaling Technology) on ice for 30 minutes, followed by washing and incubation with the Alexa fluor 488-conjugated donkey anti-mouse secondary antibody (1: 200, diluted in FACS buffer; Thermo Fisher Scientific) on ice for 20 minutes. After washing, the cells were subjected to FACS using a BD FACS AriaTM III Flow Cytometer (BD Biosciences).

Di-SG were isolated from 150-day-old porcine testes and enriched with a velocity sedimentation approach (STA-PUT), following previously published protocols (Bryant et al., 2013; Liu et al., 2015). Only fractions with high purity of Di-SG were pooled.

### Immunofluorescence

Immunofluorescence staining was performed on 4% paraformaldehyde (PFA)-fixed cytospin slides of collected cells. Briefly, the cells were permeabilized with 0.1% triton-X (Sigma-Aldrich) for 10 minutes, followed by washing and blocking with 5% bovine serum albumin (BSA; MP Biomedicals) for 1 hour. The cells were then incubated with the primary antibodies at 4°C overnight. The primary antibodies used were as follows: mouse anti-SSEA4 (1: 200; 4755S, Cell Signaling Technology), rabbit anti-UCHL1 (1: 200; ab108986, Abcam), rabbit anti-ZBTB16 (1: 200; sc-22839, Santa Cruz Biotechnology) and rabbit anti-KIT (1: 200; 3074S, Cell Signaling Technology). The corresponding isotype IgGs in place of the primary antibodies were used as negative controls. The next day, cells were washed and incubated with the Alexa fluor 488-conjugated donkey anti-mouse and/or 594-conjugated donkey anti-rabbit secondary antibodies (1: 400; Thermo Fisher Scientific) for 1 hour, followed by nuclear counterstaining with DAPI (1: 1000; Bioworld Technology) for 5 minutes. After washing, cells were visualized under a Nikon Eclipse 80i fluorescence microscope. The purity of collected spermatogonial populations was determined by the percentage of cells positive for stage-specific markers (SSEA4, ZBTB16 and UCHL1 for Un-SG and KIT for Di-SG) in the total cells (>300 cells analyzed in each group).

### Hi-C library construction

Hi-C libraries were constructed with isolated Un-SG and Di-SG, following a previously published protocol, with minor modifications (Rao et al., 2014). In brief, 1.0×10^6^ - 5.0×10^6^ cells were crosslinked with 37% formaldehyde, and then incubated with a glycine solution for 10 minutes to quench crosslinking. After washing with PBS, cells were pelleted, snap-frozen and stored at -80°C. To construct Hi-C libraries, cell pellets were resuspended in lysis buffer and homogenized. DNAs were digested with 200 units of MboI for 1 hour at 37°C. Restriction fragment overhangs were filled and labelled with biotinylated nucleotides and ligated. Ligated DNAs were then purified and sheared to 300-500bp. Ligation junctions were pulled down with streptavidin beads and subjected to Illumina NovaSeq 6000 sequencing in Novogene Co., LTD.

### Hi-C data processing

The clean Hi-C reads were mapped to the *Sscrofa* 11.1 genome and the Hi-C contact frequency between genomic loci was computed using the Juicer pipeline (version 1.8.9). Low-quality alignments (defined as MAPQ<30) and intra-fragment reads were filtered from unique reads, thereby generating valid Hi-C contacts that were used for later analyses. All contact matrices used for further analyses were KR-normalized with Juicer. The value of matrices for different samples was standardized using the R software bnbc (version 1.12.0). Correlation in contact matrices was evaluated using HiCRep (version 1.14.0), QuASAR-Rep or GenomeDISCO (Yardimci et al., 2019) in the default settings. The global interaction patterns of the whole chromosome were constructed by the scaled matrices with 100kb or 20kb bin size for independent samples. We selected 20kb to show local interactions and to perform TAD calling. To compare the high-resolution contact frequency, we merged the valid pairs from 8 independent samples of different stages and attained the KR-normalized contact matrices with the resolution of 5kb.

### Von Neumann Entropy (VNE) of intra-chromosomal contacts

The VNE was used to quantify the order in chromatin structure based on the normalized 100kb intra-chromosomal contact matrices, as previously described (Seaman & Rajapakse, 2018).

### P(s) analysis

*P*(s) analysis was performed on the normalized interaction matrices with 100kb resolution, following previously reported methods (Naumova et al., 2013). In brief, genome distances were first divided into 100kb equal bins. Then, for each bin, the mean number of interactions at corresponding distances was counted. To obtain the *P*(s), the number of interactions in each bin was divided by the total number of possible region pairs.

### RNA-seq library construction

Total RNAs were extracted from independent samples of Un-SG and Di-SG, using the RNeasy Mini Kit (Qiagen) and following the protocol provided by the manufacturer. After DNase (Qiagen) treatment, the poly A-mRNA-seq libraries were constructed with an Illumina TruSeq stranded RNA-seq library protocol.

### RNA-seq data processing

RNA-seq libraries were quantified with the Qubit dsDNA High Sensitivity Assay Kit (Thermo Fisher Scientific) and sequenced on the Hiseq 4000 platform (Illumina), producing approximately 75.75 million 150bp paired-end raw reads and 72.87 million high-quality reads for each library. Expression levels of protein-coding genes (gene annotation file [GTF] from Ensembl *Sscrofa* 11.1 release 90) were quantified as transcripts per million (TPM) using Kallisto (version 0.44.0). EdgeR (version 3.30.0) was used in differential gene expression analysis. Genes with false discovery rate (FDR) ≤ 0.05 and log_2_ (fold change) > 1 were identified as differentially expressed genes (DEGs).

### Analysis of A/B compartment

Identification of A/B compartments at 20kb resolution was performed via two steps. First, PC1 vectors were generated by using PCA as previously described at 100kb resolution (Lieberman-Aiden et al., 2009). The o/e contact matrix was then generated by the first two principal components that were obtained by using the ‘prcomp’ function in R. The initial position of gene model was defined by transcription start site (TSS) of each gene and gene density was calculated by the number of TSS in each 100kb bin. Bins with positive Pearson’s correlation between PC1 value and gene density were defined as compartment A, otherwise compartment B. Second, the A-B index, which represents the comparative likelihood of a sequence interacting with A or B, was generated as previously described at 20kb resolution (Rowley et al., 2017). Bins at 20kb resolution with the positive A-B index were considered as A compartment, and *vice versa*. The compartment strength was calculated by using AA×BB/AB^2^ as previously described (Flyamer et al., 2017). AA/BB is the mean contact enrichment between pairs of bins with compartment A/B signals, whilst AB is the mean contact enrichment between pairs of bins with compartment A and B signals. To identify genome regions that switched the A/B compartment state between Un-SG and Di-SG, the 20kb bin was defined as the A or B status in one cell type if it showed a compartment A or B signal in more than 85% of Hi-C libraries in this cell type. Genes with TSS located in A or B regions were considered as A or B genes.

### Functional enrichment analysis

Functional enrichment analysis of Gene Ontology (GO) and pathway was performed using the Metascape (http://metascape.org) (Conn et al., 2015). Genes were mapped to their human orthologs, and the lists were submitted to Metascape for enrichment analysis of the significant representation of GO Biological Process, KEGG pathway, Reactome Gene Sets and CORUM. All genes in the genome were used as the enrichment background. Cutoffs for significantly enriched terms were *P*<0.01, minimum count of 3 and an enrichment factor > 1.5. The terms were grouped into clusters based on their membership similarities.

### Analysis of TADs

Based on the normalized 20kb contact matrices, TADs were identified by employing the DI, following a previously reported method (Dixon et al., 2012b). The DI was calculated up to 2Mb flanking the center of each bin at 20kb resolution and the Hidden Markov model (HMM) was then used to predict DI states for final TAD generation. The IS for each 20kb bin was calculated as previously reported (Crane et al., 2015). The correlations of TAD architecture between samples were assessed by Jaccard indices (Stadhouders et al., 2018), and aggregate Hi-C maps were constructed as previously reported (Bonev et al., 2017). To quantify the tendency of TADs to self-interact, we calculated the D-score for each TAD, according to a previously described method (Stadhouders et al., 2018). TAD boundaries between TADs were smaller than 400kb, and the regions over 400kb were defined as unorganized chromatin. Cell type-specific TAD boundaries were identified as previously reported (Dixon et al., 2012b). To investigate TAD interaction networks, we defined TAD cliques as clusters of five or more interacting TADs in the Hi-C data, as previously reported (Paulsen et al., 2019).

### PEI identification and RP index calculation

To identify PEIs at the resolution of 5kb, we generated aggregated Hi-C maps for each cell type. PEIs were identified by applying PSYCHIC based on the 5kb contact matrices (Ron, Globerson, Moran, & Kaplan, 2017). The genome was divided into TADs and similar neighboring domains were further merged into a hierarchical structure. Then, a domain-specific background model was built according to the fitted bilinear power-law model for each or merged TADs. High-confidence PEIs were identified by interaction intensity normalized by the background model with FDR value < 0.001 and interaction distance ≥ 15kb.

Later, we calculated the RP index that combines both the number and intensity of the interacting enhancers to quantify the potential of PEIs for transcriptional regulation of target genes. RP=Σn[log_10_ (normalized interaction intensity of PEIs)], where n refers to the number of interacting enhancers. The normalized interaction intensity of PEIs was calculated by the observed contact frequency minus the background contact frequency.

### ChIP-seq library construction

The ChIP assay was conducted as previously described (Han, Ren, Cao, Zhao, & Yu, 2019). In brief, 1.0×10^6^ - 5.0×10^6^ cells were crosslinked with 37% formaldehyde, and then incubated with a glycine solution for 10 minutes to quench crosslinking. After washing with PBS, cells were pelleted and lysed. Chromatins were sonicated to obtain the sheared 200-500bp DNA. Around 20uL chromatin was saved as input DNA at -20°C, and 100uL chromatin was used for IP with the H3K4me3 antibody (9751, Cell Signaling Technology) or the H3K27ac antibody (ab4729, Abcam). Approximately 5μg antibody was used in each IP reaction at 4°C overnight. The next day, 30μL protein A/G beads were added and subjected to further incubation for 3 hours. After washing, the binding materials were eluted from the beads. The immunoprecipitated DNAs were then used to construct the ChIP-seq library, following the protocol provided by the manufacturer and sequenced on Illumina Xten with the PE 150 method.

### ChIP-seq data processing

Trimmomatic (version 0.38) was employed to filter out low-quality reads (Bolger, Lohse, & Usadel, 2014). The cleaned ChIP-seq reads were aligned to the pig genome (*Sscrofa* 11.1), using the BWA (version 0.7.15) with default settings. The independent samples of each cell type were pooled using SAMtools (version 0.1.19). To identify enriched regions of active markers (H3K4me3 and H3K27ac), peak calling was performed using SICER (version 1.1).

### Annotation of PEIs with ChIP-seq data

To define active enhancer-involved PEIs, we first identified REs and SEs by using the ROSE algorithm (Loven et al., 2013; Whyte et al., 2013). Next, we divided all SEs into two categories as previously reported, to which we referred as hierarchical and non-hierarchical SEs (Huang et al., 2018). The hierarchical SEs were then divided into hub and non-hub enhancers by applying a threshold value of H-score which corresponds to the 90th percentile of Z-score. The active enhancer-involved PEIs were defined if the 5kb enhancer bin overlapped with the identified enhancer by at least 1bp.

### Statistics

Statistical comparisons were conducted by using the Mann-Whitney U test (one-tailed), unless otherwise stated. A difference was considered significant when *P*<0.05.

## Data availability

The raw and processed data sets generated in this study will be available in the NCBI SRA database under the accession number PRJNA743697.

## Author contributions

Y.Z., L.Z., L.J., P.Z., M.L. and W.Z. conceived the study; Y.Z., L.Z., L.J., P.Z., F.L., M.G., Q.G. and Y.Z. collected the data; Y.Z., L.Z., L.J., P.Z. and M.L. performed the analyses; Y.Z. and L.Z. drafted the original manuscript; Y.Z. and L.J. revised the manuscript; M.L. and W.Z. supervised the study. All authors read and approved the final version and submission.

## Supplementary files

**Fig S1.**
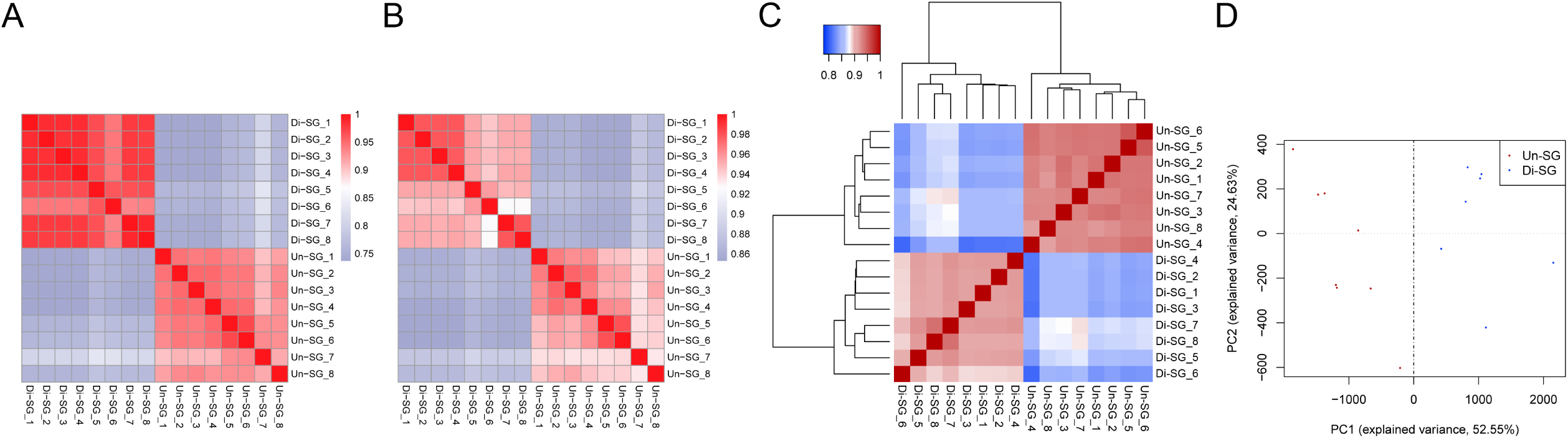
The correlation of normalized Hi-C interaction matrices between Un-SG and Di-SG samples illustrated by QuASAR-Rep (A), GenomeDISCO (B), Pearson correlation analysis (C) and PCA plot (D).

**Fig S2.**
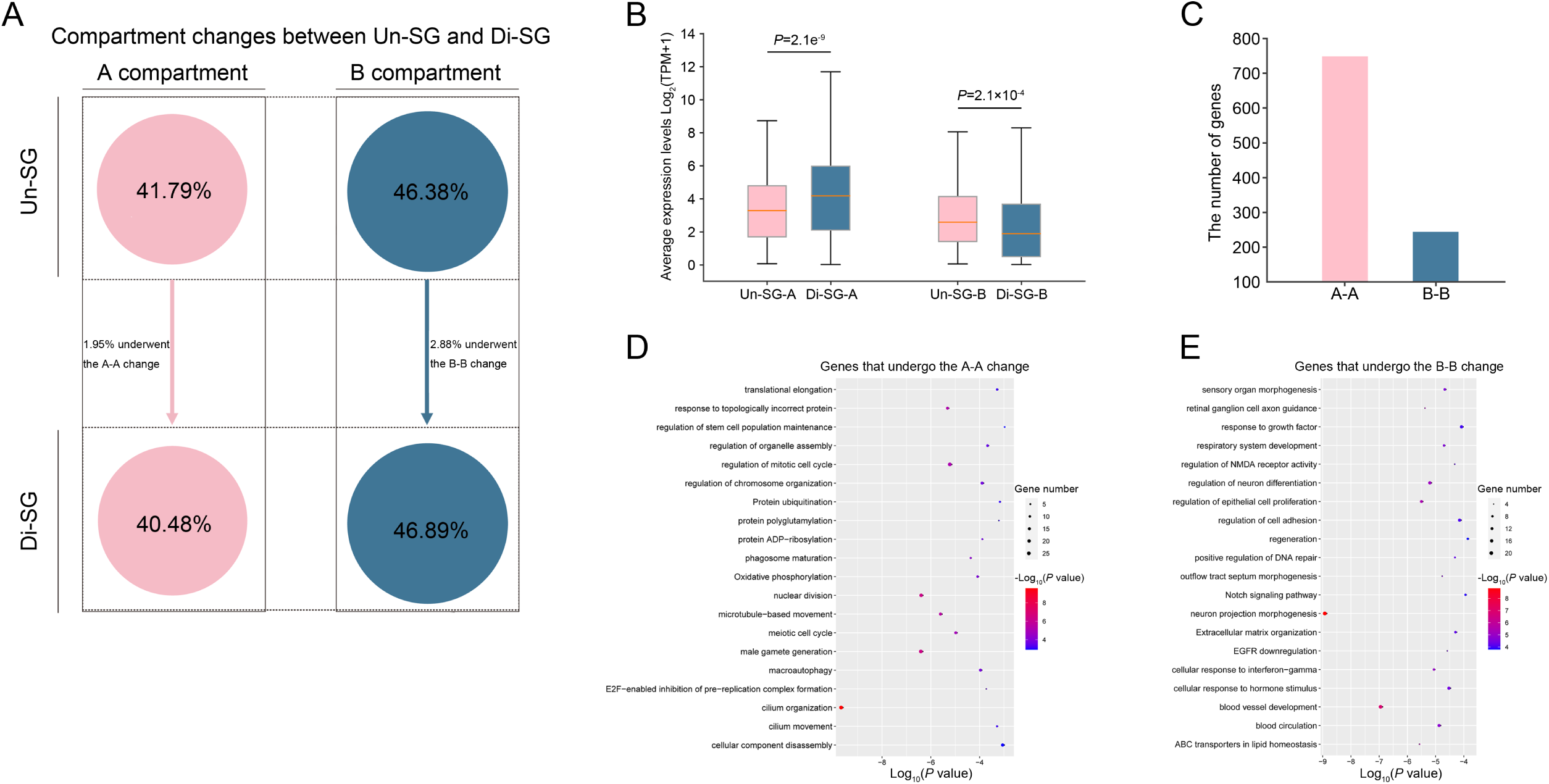
A/B compartment changes during spermatogonial differentiation. (A) A schematic overview illustrating the proportions of genomic regions subjected to A/B compartment changes (A-A or B-B) between Un-SG and Di-SG. (B) The average expression levels of genes that underwent the A-A or B-B change. *P*: Mann-Whitney U test, one-tailed. (C) The numbers of genes that underwent the A-A or B-B change. (D and E) GO-BP analysis of genes that underwent the A-A (D) or B-B change (E).

**Fig S3.**
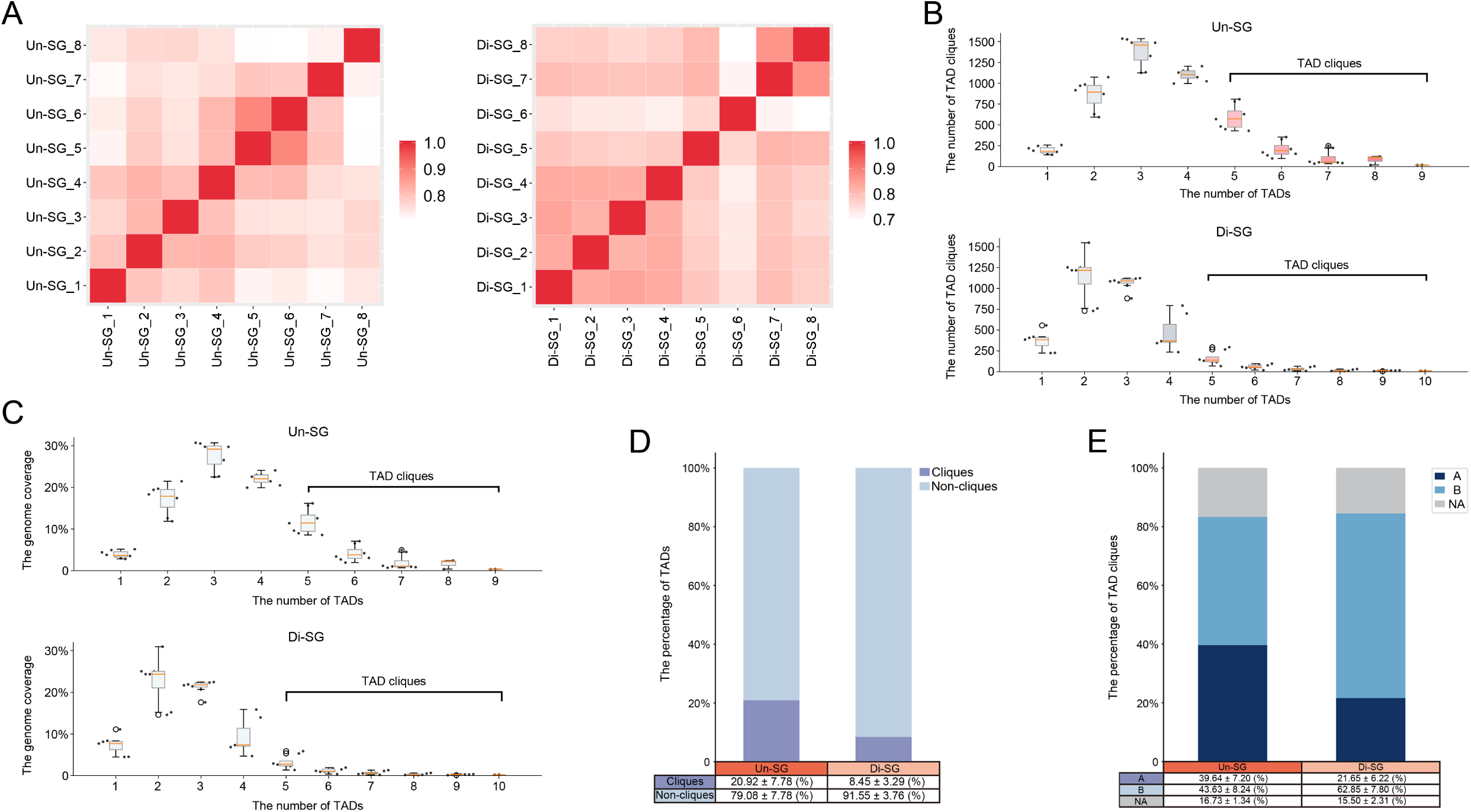
TAD cliques in spermatogonial subgroups. (A) Pearson correlation analysis illustrating the correlation of TAD cliques between Un-SG (left) and Di-SG (right) ingroup samples. (B) The numbers of TAD cliques in Un-SG and Di-SG. Data points refer to eight independent samples. (C) The genome coverage by TAD cliques in Un-SG and Di-SG. Data points refer to eight independent samples. (D) The proportions of TADs in cliques and non-cliques. Data are presented as the mean ± SD of eight independent samples. (E) The proportions of TAD cliques in A/B compartments. Data are presented as the mean ± SD of eight independent samples.

**Fig S4.**
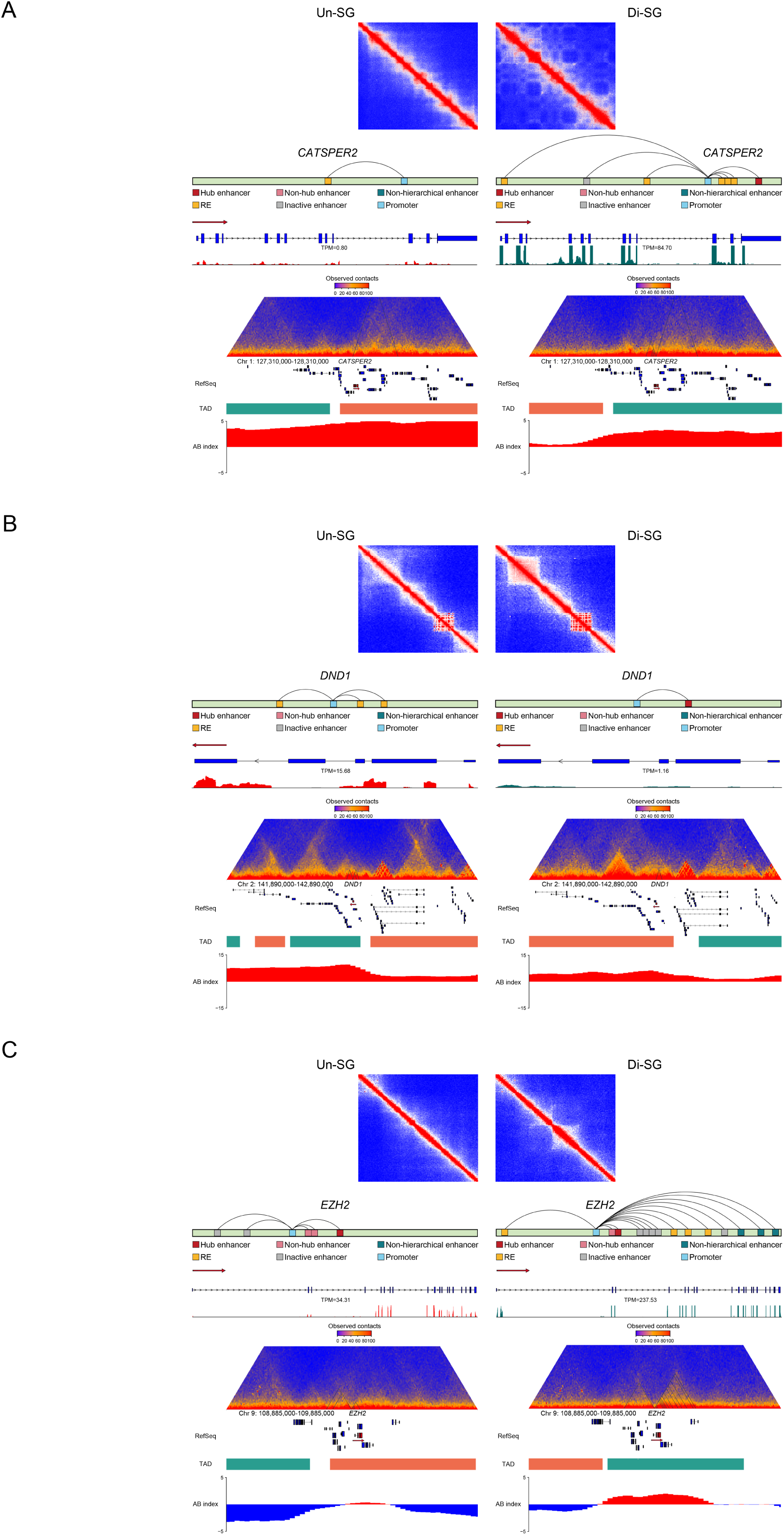
PEI regulation of *CATSPER2* (A), *DND1* (B) and *EZH2* (C) in Un-SG and Di-SG. The upper panel: the contact matrix showing stripes at the *CATSPER2* (A), *DND1* (B) and *EZH2* (C) SE loci. The middle panel: models for PEI regulation of *CATSPER2* (A), *DND1* (B) and *EZH2* (C), as well as RNA-seq coverage at their genomic loci. The lower panel: views of the observed/expected chromatin interaction frequencies, RefSeq, TAD and A/B index at chromosome 1: 127,310,000-128,310,000 (A), chromosome 2: 141,890,000-142,890,000 (B) and chromosome 9: 108,885,000-109,885,000 (C).

**Table S1**. Hi-C and RNA-seq data summaries.

**Table S2**. List of genes subjected to the A-B/B-A/A-A/B-B change.

**Table S3**. GO-BP analysis of genes subjected to the A-B/B-A/A-A/B-B change.

**Table S4**. List of genes harbored in Un-SG- or Di-SG-specific TAD boundaries.

**Table S5**. GO-BP analysis of genes harbored in Un-SG- or Di-SG-specific TAD boundaries.

**Table S6**. List of genes with differential RP indices.

**Table S7**. GO-BP analysis of genes with differential RP indices.

**Table S8**. List of genes with Un-SG- or Di-SG-exclusive PEI regulation.

**Table S9**. List of RE- and hierarchically organized SE-associated genes.

**Table S10**. GO-BP analysis of RE and SE-regulated genes with differential RP indices and expression levels.

